# Analysis of sea star larval regeneration reveals conserved processes of whole-body regeneration across the metazoa

**DOI:** 10.1101/118232

**Authors:** Gregory Cary, Andrew Wolff, Olga, Joseph Pattinato, Veronica Hinman

## Abstract

**Background:** Metazoan lineages exhibit a wide range of regenerative capabilities that vary among developmental stage and tissue type. The most robust regenerative abilities are apparent in the phyla Cnidaria, Platyhelminthes, and Echinodermata, whose members are capable of whole-body regeneration (WBR). This phenomenon has been well-characterized in planarian and hydra models, but the molecular details of WBR are less established within echinoderms, or any other deuterostome system. Thus, it is not clear to what degree aspects of this regenerative ability are due to deeply conserved mechanisms.

**Results:** We characterize regeneration in the larval stage of the Bat Star (*Patiria miniata*). Following bisection along the anterior-posterior axis, larvae progress through phases of wound healing and re-proportioning of larval tissues. The overall number of proliferating cells is reduced following bisection and we find evidence for a re-deployment of genes with known roles in embryonic axial patterning. Following axial re-specification, we observe a significant localization of proliferating cells to the wound region. Analyses of transcriptome data highlight the molecular signatures of functions that are common to regeneration, including specific signaling pathways and cell cycle controls. Notably, we find evidence for temporal conservation among orthologous genes involved in regeneration from published Platyhelminth and Cnidarian regeneration datasets.

**Conclusions:** These analyses show that sea star larval regeneration includes phases of wound response, axis respecification, and wound proximal proliferation. Commonalities of the overall process of regeneration, as well as gene usage between this deuterostome and other species with divergent evolutionary origins suggest a deep conservation of whole-body regeneration among the metazoa.

## BACKGROUND

The evolution of regenerative abilities has fascinated researchers for centuries. Species with a capacity for restorative regeneration are distributed throughout the metazoan tree of life (Figure 1A), however the extent to which any animal is capable of regenerating varies considerably. Whereas some taxa are able to undergo whole-body regeneration (WBR), other lineages exhibit much more restricted regenerative capabilities (*e.g.,* the ability to re-grow only specific organs or tissues) [1–3]. Given the broad phylogenetic distribution of robust regenerative abilities, it remains unclear if elements of this phenomenon are directed by deeply conserved molecular mechanisms that have been lost in species with more restricted regenerative capacities or have evolved multiple times independently. While many attempts have been made to synthesize regenerative phenomena in disparate taxa [1–3], or to provide evolutionary context to genes utilized during regeneration within a particular model [4, 5], few studies have directly compared the transcriptional control of regeneration among highly regenerative, distantly-related metazoan lineages. This is, in part, because we are as yet missing detailed descriptions of regeneration from key taxa. By approaching regeneration from an evolutionary perspective, it is possible to identify conserved mechanisms that underlie regenerative abilities. This has significant implications for if and how regeneration can be induced in organisms with more limited potential.

**Figure 1.**
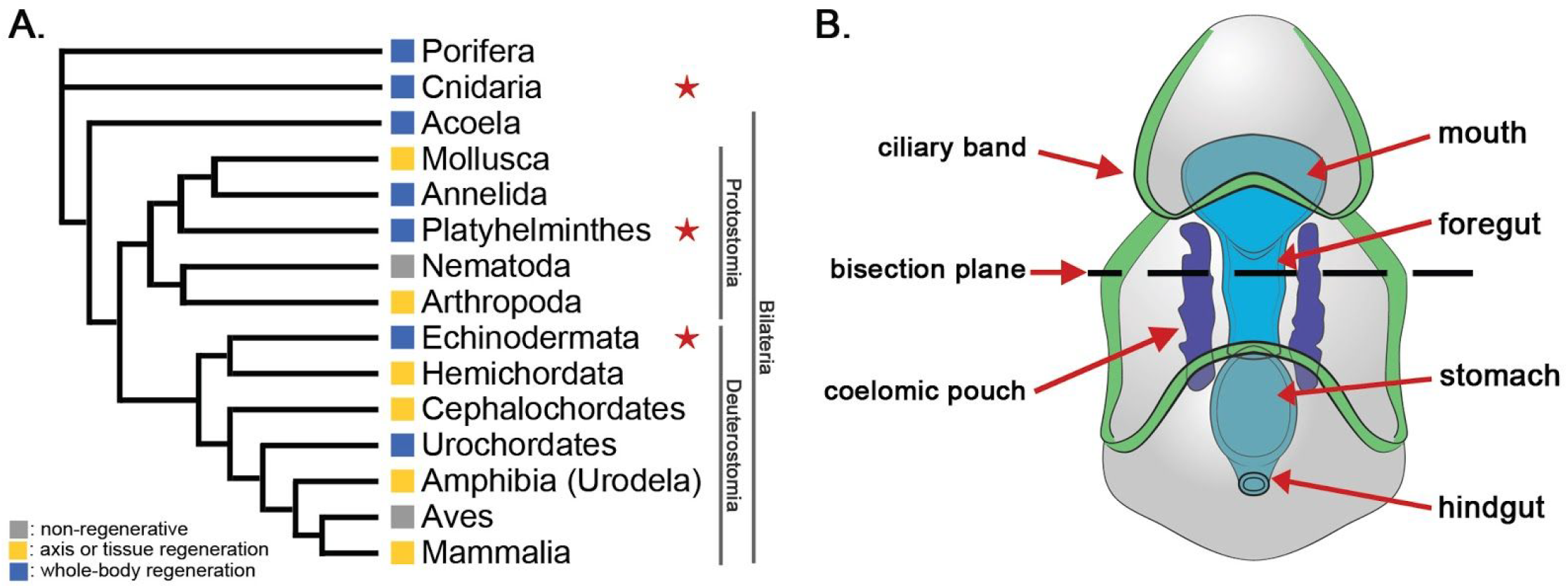
Models of whole-body regeneration. (A) Phylogeny depicting regeneration capacity of various taxa, after [2, 88]. Species from the three taxa marked with a star were considered in this study. (B) Schematic of a sea star bipinnaria larva indicating the bisection plane (dashed line) and relevant anatomical features including the ciliary band epithelium (green), coelomic pouch epithelium (purple), and enteric organs (blue).

The best characterized models for understanding regeneration are species of Cnidaria (*e.g. Hydra vulgaris [6, 7]*) and Planaria (*e.g. Schmidtea mediterranea [8, 9]*). These organisms are capable of WBR, meaning that they can regrow all body parts following amputation [2]. In these contexts, WBR involves transitions through wound healing, immune signaling, axis/organizer specification (especially via WNT signaling), and differentiation of new cells to replace missing cells and tissues [7–11]. A key distinction between these models lies in the source of the newly differentiated cells. In planarians (bilaterian protostomes within the phylum Platyhelminthes), a pool of somatic stem cells (neoblasts) generates a proliferative blastema that is essential for regeneration [12–14]. In contrast, regeneration in *Hydra* species is mediated through de-differentiation and transdifferentiation of existing cells to replace those lost by injury [15, 16], in addition to somatic stem cells (interstitial cells or I-cells), which serve as both undifferentiated precursors of several cell types [17] and also proliferate following injury [18].

Regenerative ability is generally more limited in deuterostomes. Within vertebrates, regeneration is frequently restricted to specific developmental stages, tissues, or organs [2]. By contrast, many invertebrate deuterostomes are capable of extensive regeneration of all tissues at multiple developmental stages. Colonial ascidians (*e.g. Botryllus schlosseri*) are capable of WBR [19, 20], whereas solitary species are capable of partial regeneration (*e.g.* adult siphons in *Ciona intestinalis*) [21, 22]. Hemichordate species (*e.g. Ptychodera flava*) can regenerate the adult head when bisected from the body [23, 24]. However, the best known and most regenerative species of deuterostomes belong to the Echinodermata.

Echinoderms (*e.g.* sea stars, brittle stars, and sea cucumbers) exhibit remarkably robust regenerative capabilities throughout all life stages. Adult echinoderms have been a focus of regeneration studies that examined re-growth of specific structures (*e.g.* spines, tube feet, nerve cord, gut, and arms) [25–39]. Regeneration has also been observed in larvae from all echinoderm classes examined [40]. These planktonic stage echinoderms can swim and feed in the water column for weeks or months. Larval regeneration is more similar to the WBR observed in planaria and hydra, as it requires the complete re-growth of all tissues and organ systems. Molecular studies of regenerating sea star larvae have identified several regeneration-specific changes in gene expression, including the *sea star regeneration-associated protease* (SRAP; [41]), *vasa*, *nodal*, *dysferlin*, and *vitellogenins* (*vtg1* and *vtg2*) [42]. However, to date, a comprehensive survey of gene expression changes during larval echinoderm regeneration has not been reported. As one of the few deuterostome taxa capable of undergoing WBR, sea star larvae can provide unique insight into the evolution of regenerative processes.

Here, we characterize the molecular and cellular events that occur during regeneration in the larval asteroid *Patiria miniata* and assess the expression patterns of orthologous genes in other distantly related species that undergo WBR. We first characterize the landmark regeneration events: wound healing, tissue re-proportioning, cellular proliferation, and cell death. To characterize the transcriptional changes that underpin these events, bisected larval fragments were evaluated using RNA-Seq. Through analysis of these data we define broad gene classes that are expressed similarly in both anterior and posterior regenerating fragments. Finally, through identification of orthologous genes between *P. miniata* and published datasets of regenerating hydra and planarian models (Figure 1A), we find sets of genes that have similar temporal expression profiles in these distantly-related regenerating organisms. These results highlight similarities in the regeneration programs of a bilaterian deuterostome, a lophotrochozoan, and a basally branching eumetazoan. This suggests that regeneration may be common to the base of all animals.

## RESULTS & DISCUSSION

### Bipinnaria regeneration involves wound healing, body re-proportioning, cell proliferation and cell death

To make an informed comparison to other regenerative models, we first characterized the stages of larval regeneration in *P. miniata*. Bipinnaria larvae (7 days post-fertilization [dpf]) were bisected midway along the transverse Anterior-Posterior (AP) axis (Figure 1B). Both resulting larval fragments were completely regenerative, restoring all lost tissues and organs over the course of two weeks. These findings are consistent with previous reports of larval asteroid regeneration [42, 43]. Although we focus on the regeneration of the posterior fragments, a similar regenerative response is apparent within the anterior fragment (Figure 2-S2).

**Figure 2.**
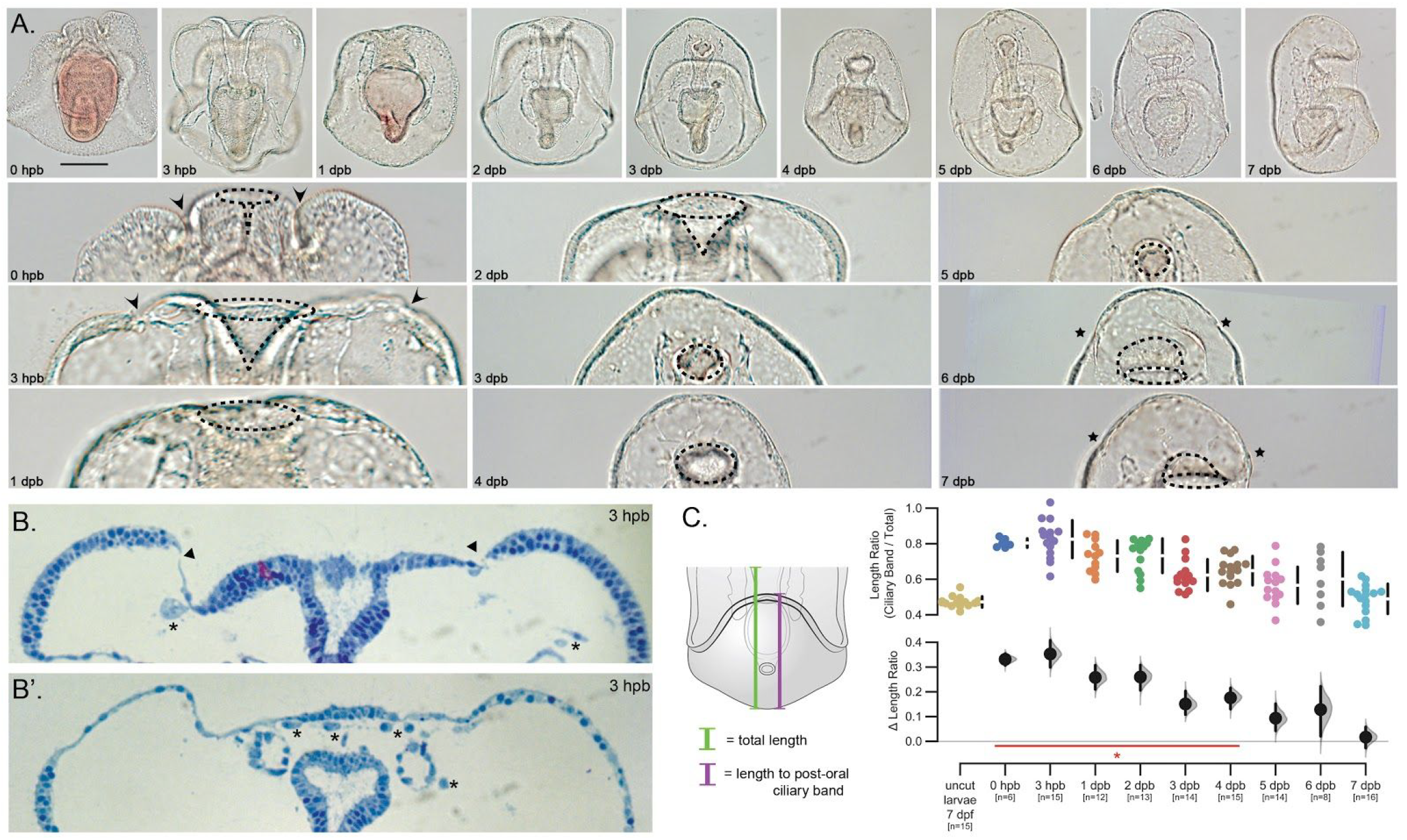
Sea star bipinnaria regeneration involves wound healing, re-proportioning and re-specification. (A) DIC images showing larval recovery following bisection (top row) and magnifications of the wound site at each stage (bottom row). Important anatomical features are highlighted in the magnified images including the wound site (arrowheads), opening to the gut lumen (dotted lines), and new ciliary bands (asterisks). Scale bar = 100 μm; applicable to all images in panel. (B) Two serial sections from the same individual showing wound closure (arrowheads) and many free cells within the blastocoelar space (asterisks). (C) Ratios of length from the posterior pole to the top of the post-oral ciliary band to length from the posterior pole to the anterior pole (i.e. total length of the specimen) are plotted along with the difference of the means (i.e. Δ Length Ratio) and 95% confidence interval. Those timepoints with a ratio found to be significantly different than uncut larvae are indicated by the red line and asterisk (Mann-Whitney U test, p-value < 0.001). n=number of individuals measured at each timepoint.

We observe that the initial wound is mostly closed by 3 hours post-bisection (hpb; Figure 2A-B, arrowheads). This also coincides with the appearance of several types of mesenchymal blastocoelar cells proximal to the wound epithelium. After this rapid wound healing response, larvae re-proportion their remaining tissues over the first several days post-bisection (dpb). This is evident when analyzing the position of the post-oral (lower) ciliary band (Figure 2C). Prior to bisection, this ciliary bandis located in the middle of the larva; on average, the distance from the posterior end of the larva to the ciliary band is 47% of the total length of the larva (Figure 2C). Immediately after bisection, this ratio increases to 80% as the anterior region has been removed (Figure 2-S1). However, over the subsequent five days the larval proportions return to pre-bisection ratios (at 5 dpb, the ciliary band to larval length ratio is 57%). Importantly, this reallocation of tissues is not due to an increase in the total length of the larval fragments, as we show that the overall length of the bisected larva does not change during this time (Figure 2-S1). Although we did not quantify the change, we note a similar re-proportioning of the larval midgut between 1 dpb and 5 dpb, and also observed that the shape and position of the larval mouth changes. During bisection, the foregut is cut in half such that the anterior portion forms a new oral opening oriented along the anterior-posterior axis. However, by 3 dpb the oral opening is reoriented ventrally and tissues are apparent anterior to this opening. Finally, by 6 dpb we observe the return of most morphological features, including the anterior ciliary band, the oral field and oral lobe. Together, these findings indicate that regeneration in larval sea stars occurs in at least three stages: healing at the wound site, tissue reallocation, and restoration of lost tissue. Similar patterns are evident in regenerating anterior fragments (Figure 2-S2).

We next analyzed the pattern of cellular proliferation during regeneration. Larvae were exposed to EdU (6 hr pulses) to mark proliferating cells in normal (uncut) and over the course of larval regeneration (Figure 3). In uncut larvae, EdU^+^ cells are widely distributed (Figure 3A). We infer from this result that larvae are actively growing. However, upon bisection, the numbers of EdU^+^ cells steadily decrease (Figure 3B; Mann-Whitney P < 2×10^−4^). This decrease in EdU^+^ cell number is accompanied by a change in the localization of proliferating cells. EdU^+^ cells localize proximal to the wound sites (3 dpb in posterior fragments and 6 dpb in anterior fragments) and fewer EdU^+^ cells are located in more distal tissues distal (Figure 3C; Mann-Whitney P < 0.05). Moreover, the proliferating cells that localize to the wound site are distinct from cells that proliferate early. Cells proliferating at 1 dpb were labeled with pulse of BrdU followed by a wash-out. Cells proliferating during the later phases were then labeled with a pulse of EdU, and processed for imaging. We find very little overlap of BrdU^+^ cells that are also EdU^+^ (Figure 3D). This indicates that cells proliferating during early regeneration do not to continue to divide during the later, wound-proximal proliferation phase of regeneration. In non-bisected, stage equivalent control larvae, by contrast, there is extensive overlap between BrdU^+^ and EdU^+^ cells (Figure 3D). This suggest that under normal conditions, cells that are proliferating normally continue to divide, but following bisection, different populations of cells now enter proliferation. Thus, during the regenerative response, typical, system-wide larval growth is inhibited, and regeneration-specific cell proliferation is concentrated at the regenerating edge where tissues later form.

**Figure 3.**
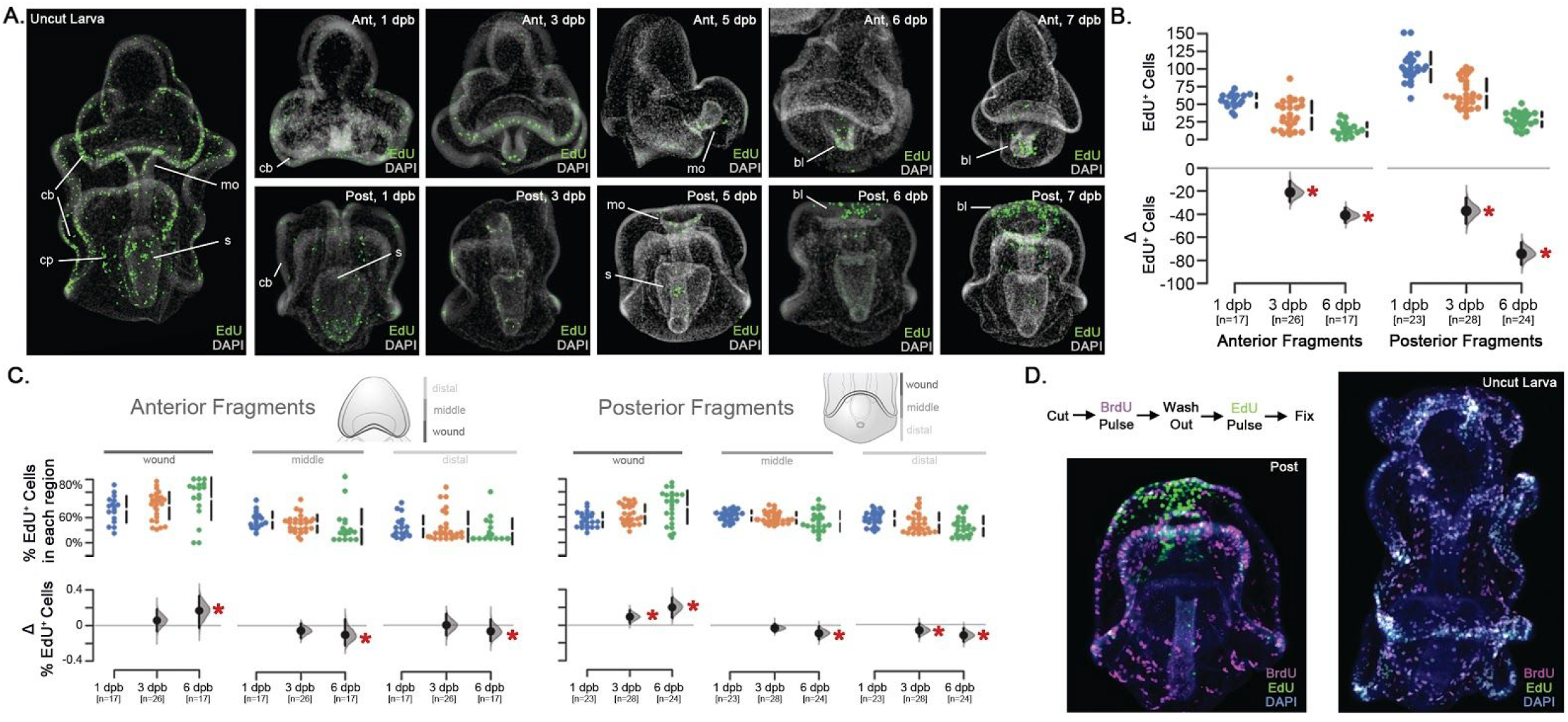
Cell proliferation decreases and localizes to wound-proximal cells. (A) EdU staining of S-phase cells in intact and regenerating sea star larvae (1-7 days post bisection, dpb). EdU positive cells are shown in green. Nuclei were stained with DAPI and shown in gray. Cell proliferation in uncut larvae is throughout the ciliary band epithelium (cb), mouth (mo), stomach (s), and coelomic pouches (cp). Regenerating anterior fragments (top row) and posterior fragments (bottom row) demonstrate similar initial distributions of proliferation, although the number of EdU^+^ cells decreased by 3 dpb. Beginning at 6 dpb, EdU^+^ cells are concentrated near the wound site in both anterior and posterior regenerating fragments in a putative regeneration blastema (bl). (B) Quantitation of the EdU^+^ cells shows a steady decline in the number of proliferating cells in both anterior and posterior regenerating fragments. The difference of the means (i.e. Δ EdU^+^ Cells) is plotted and significance differences are indicated (Mann-Whitney, p < 0.05, red asterisk). n= total number of bisected animals counted. (C) The fraction of EdU^+^ cells in each of the wound-proximal, middle, and wound-distal thirds of each regenerating larval fragment from panel B is shown. The number of individuals counted is the same as in (B). The difference of the means (i.e. Δ % EdU^+^ Cells) is plotted and significance differences are indicated (Mann-Whitney, p < 0.05, red asterisk). (D) The experimental regimen of the BrdU/EdU pulse-chase experiments is shown. Regenerating larvae (left) or uncut larvae (right) were labeled with BrdU (magenta) for 6 hours after which the BrdU was washed out. Larvae are subsequently labeled with a 6 hour EdU pulse (green) at the onset of wound-proximal proliferation, or after a similar duration for uncut larvae.

As a corollary to understanding cell division during larval regeneration, we examined the patterns of cell death using TUNEL assays. In normal larvae, TUNEL^+^ cells are distributed organism-wide (Figure 4A). Following bisection, the number and distribution of apoptotic cells remains largely unchanged for several days (Figure 4B-D and 4-S1). However, at 6 dpb there is a significant increase in the total number of TUNEL^+^ cells in both anterior and posterior regenerating fragments (Mann-Whitney P < 4×10^−5^). Unlike cell proliferation, these cells are not preferentially located with respect to the wound epithelium (Figure 4-S1B). Together, these results indicate that regeneration induces a global decrease in cell proliferation, followed by a rapid increase in cells cycling near the wound site. In contrast, the rate of cell death is consistent and increases across the larva coincident with the onset of wound-localized cellular proliferation.

**Figure 4.**
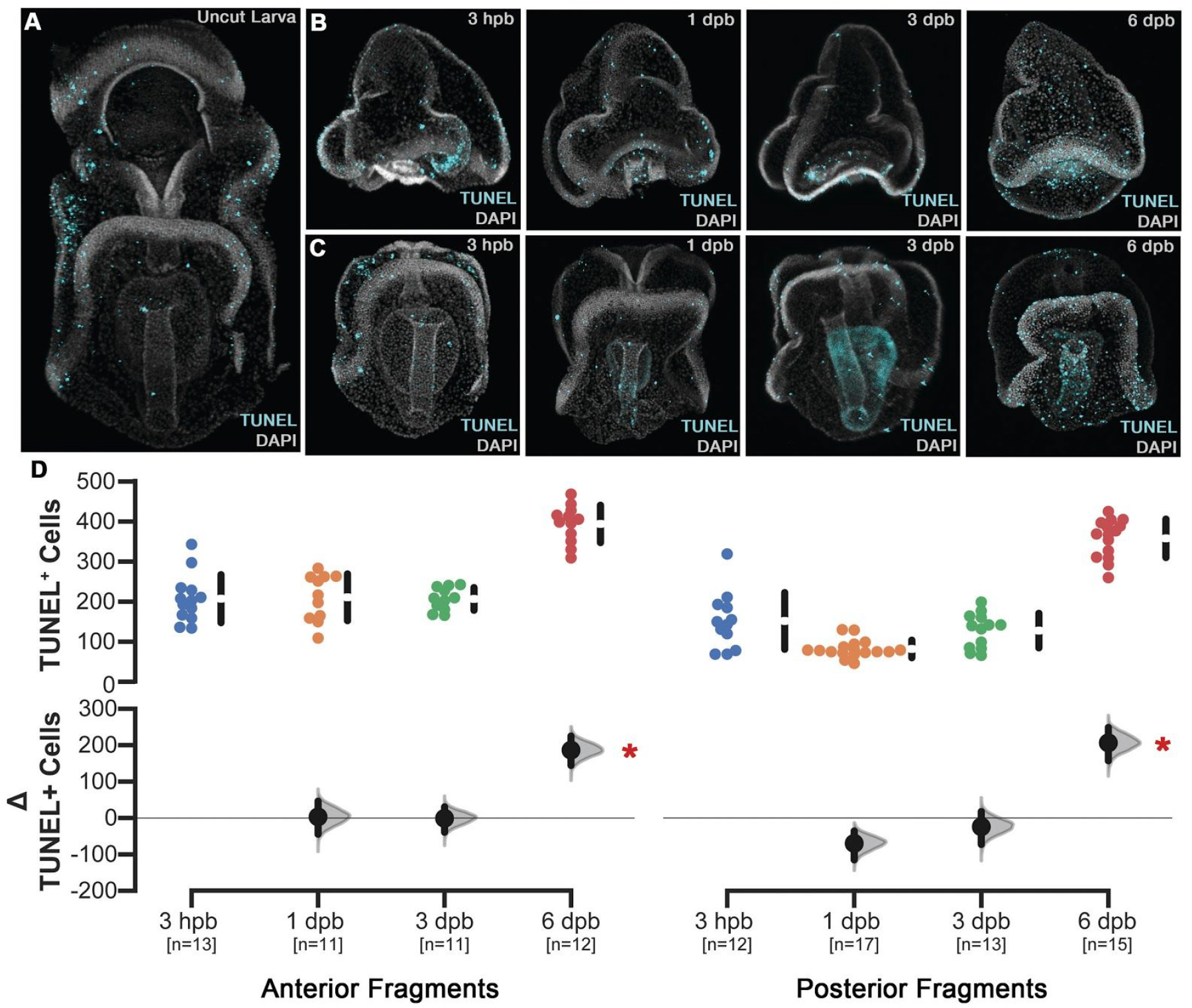
Apoptotic cell death persists and increases in later phases. (A) TUNEL^+^ cells (green) in control animals are normally distributed throughout larval tissues and is concentrated within the ciliary band epithelium. Nuclei (gray) stained with DAPI. Regenerating anterior (B) and posterior (C) fragments display similar patterns and numbers of TUNEL^+^ cells from 3 hours post bisection (hpb) until 6 days post bisection (dpb) when there is an increase. (D) Quantitation of TUNEL^+^ cells in regenerating anterior and posterior fragments shows that there is no significant difference in the number of TUNEL^+^ cells until 6 dpb when a significant increase in apoptotic cells are detected. The difference of the means (i.e. Δ TUNEL^+^ Cells) is plotted and significance differences are indicated (Mann-Whitney, p < 3×10^−4^, red asterisk). n= the number of individuals sampled.

These cellular and tissue changes during larval sea star regeneration define landmark features of the regenerative process including wound healing, re-proportioning of larval tissues, and onset of wound-proximal proliferation along with a coincident increase in apoptotic cell death. These broad characterizations mirror regenerative processes described in other organisms, and suggest a shared toolkit of regenerative responses.

### Transcriptome analyses of larval regeneration explain the genetic basis underlying observed cellular and morphological phenomena

To characterize the molecular events that operate during larval sea star regeneration and to establish a dataset amenable to inter-species comparison, we surveyed gene expression changes across a time course of larval regeneration. Pools of regenerating posterior fragments, anterior fragments, and non-bisected sibling control larvae were collected at three points following bisection: one early time point (approximately 3 hpb), one intermediate time point (3 days post bisection, dpb), and one time point at the initiation of wound-localized cell proliferation (6 dpb). By separately sampling RNA from each pool of regenerating fragments, we were able to identify changes in gene expression changes that occur in both the anterior and posterior fragments as well as those that are specific to regeneration in each context. The inclusion of non-bisected, age-matched, sibling larvae control for transcriptional changes due to continuing larval development as well as genetic differences among cultures. For each time point, transcript levels were compared between each pool of regenerating fragments and the control larvae (*i.e.,* anterior vs. uncut and posterior vs. uncut). In total, 9,211 differentially expressed genes (DEG) were identified from these comparisons.

We implemented a hierarchical clustering approach to distinguish fragment-specific expression patterns from expression changes that are shared in both regenerating fragments (Figures 5A and 5-S1). In total, five expression clusters were identified: (I) genes upregulated early in both anterior and posterior fragments; (II) genes downregulated early in both fragments; (III) genes up in the anterior, down in posterior; (IV) genes up in the posterior, down in anterior; and (V) genes up-regulated later (*i.e.* by 6 dpb) in both fragments (Figure 5A). Thus, we have identified three subsets of DEGs that exhibit similar expression profiles during regeneration in both fragments (*i.e.* Clusters I, II, and V) and two subsets that are strongly fragment-specific (*i.e.* Clusters III and IV). To validate the RNA-seq measurements, we analyzed the same samples using a custom Nanostring nCounter codeset. In total 69 of the 74 genes (92.3%) tested by our Nanostring experiments exhibited either a similar trend and significance status or just a similar trend to the measurements made by RNA-seq (Figure 5-S2).

**Figure 5.**
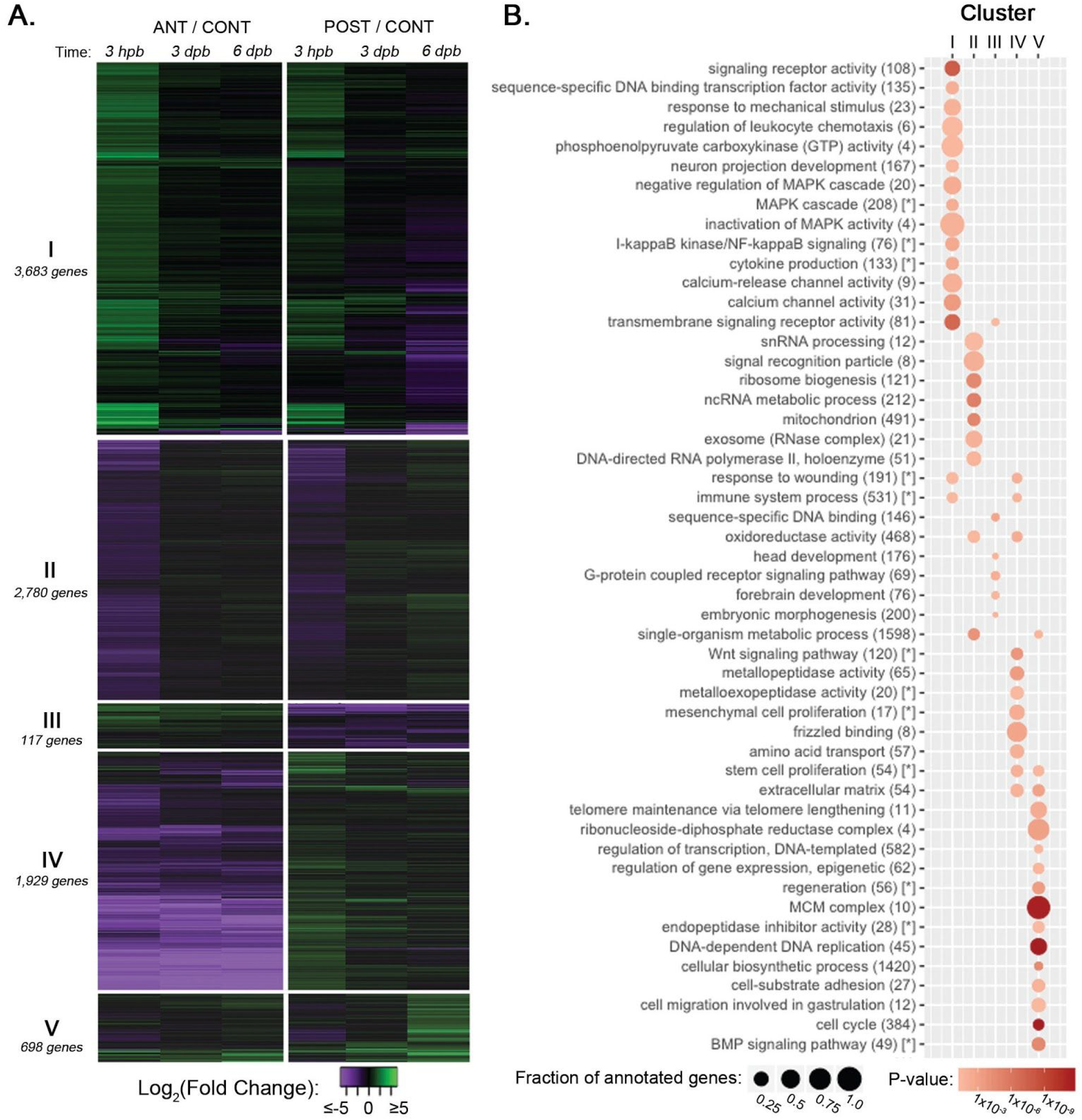
Cluster analysis indicates genes involved in regenerative functions. (A) The heatmap depicts log fold change values for genes (rows) in anterior (ANT) and posterior (POST) regenerating fragments compared with sibling uncut control larvae (CONT) over the sampled regeneration time points (columns; 3 hours post bisection hpb, 3 days post bisection, dpb, and 6 dpb). Green indicates a positive fold change (upregulated with respect to uncut controls), whereas purple indicates a negative fold change (down-regulated with respect to control). (B) Gene ontology (GO) term enrichments for each of the five clusters. The enrichment of each GO term is indicated by a circle where the area corresponds to the fraction of genes annotated with that term are present in the cluster, and the color of the circle corresponds to the corrected hypergeometric p-value of term enrichment. Terms marked with a [*] are from the annotation set generated by mouse gene ortholog prediction (Figure 5-S3).

To provide further insight into the functions of genes that were assigned to each cluster, we identified enriched Gene Ontology (GO) terms (Figures 5B and 5-S3). Genes in Clusters I and II (*i.e*. genes that are up-or down-regulated early in both regenerating fragments), are enriched for GO terms associated with a robust wound response. Up-regulated genes (Cluster I) are enriched for terms that include cell signaling pathways (*e.g.* “MAPK cascade” and “calcium channel activity”), “response to wounding”, and “immune system process” (Figures 5B and 5-S3). This cluster is also enriched for terms that indicate an early involvement of innervation and ciliogenesis (*e.g.* “neuron projection development” and “motile cilium”) which are common in other regeneration models [44–47]. The down-regulated genes (Cluster II) are enriched for terms that point to a shut-down of anabolic processes (“ribosome biogenesis”, “gene expression”) as well as primary metabolism (e.g. “mitochondrion” and “metabolic process”). Together, these clusters of early-regulated genes are consistent with a rapid response to the bisection insult that involves down-regulation of highly energetic cellular processes and up-regulation of functions that are specific to the injury response.

Clusters III and IV are composed of genes whose profiles are highly fragment specific; these genes are differentially regulated in each fragment relative to control larvae. Many of these genes are expressed asymmetrically along the AP axis. Thus, bisection results in the loss of posterior-specific gene expression from anterior fragments and vice versa. For example, Cluster III is enriched for genes annotated with functions specific to anterior larval fragments, such as “head development” [48], whereas Cluster IV is enriched for genes associated with posterior fates in embryonic sea stars, such as “Wnt signalling pathway” [49].

Finally, although Cluster V is comprised of relatively few genes, it is the most functionally coherent cluster. That is, the GO term enrichment analyses are the most statistically significant and reproducible across the three sources of functional annotations tested (Figure 5B and 5-S3). Genes assigned to Cluster V are enriched for terms related to the cell cycle, DNA replication, and extracellular matrix (ECM) remodeling. The Cluster V genes, which are up-regulated late (by 6 dpb) in both fragments, likely reflect the onset of localized cellular proliferation that occurs at this time (Figure 3A). Importantly, these genes are up-regulated in regenerating fragments although the total number of proliferating cells has decreased compared to controls (Figure 3A). This suggests that the Cluster V genes represent a regeneration-specific increase in expression of proliferation associated genes that is distinct from the normal, growth-associated proliferation.

### Comparative transcriptome analyses reveal homologous genes with shared expression profiles among distantly related animals

Having identified the overall morphological progression of larval sea star regeneration (*i.e.* wound response, axis re-proportioning, and cell proliferation), we sought to determine if orthologous genes with similar temporal expression exist in other models of WBR. Such homology could indicate not only a shared overall progression, but that the genes involved are also conserved. To address this question, we used published transcriptome data from regenerating planaria (*S. mediterranea*) [4] and hydra (*H. magnipapillata*) [5] for comparison. The Kao *et al.* dataset [4] was selected because it consolidated several planarian transcriptome assemblies, resulting in a more complete gene set, and also independently sampled both regenerating anterior and posterior worms, which is analogous to our own study design. Furthermore, the time points sampled range from 0 hours post-amputation (hpa) to 72 hpa, at which point planarian blastemal proliferation reaches its peak [9]. This time frame roughly corresponds to the phases of regeneration considered in our study of larval sea stars. Regeneration has been less well-characterized from a molecular standpoint in hydroids; the Petersen *et al.* dataset [5] is the only available transcriptome study from regenerating hydra. Here, RNA was sampled only from the distal tip of regenerating aboral tissues during the 48 hours it takes to achieve complete head regeneration. As blastemal proliferation is not a feature of hydra regeneration, this characteristic cannot be used to synchronize the regenerative phases in this study to the other datasets. Nonetheless, these published datasets provide the best available basis for comparison to our sea star dataset.

To identify orthologs that share similar temporal dynamics during regeneration, the reported expression values from each dataset were clustered. For each comparative dataset, we assigned genes to three coarse clusters: those that were up-regulated early in regeneration and down-regulated later, those that were downregulated early in regeneration and up-regulated in later regeneration, and those that exhibited some other temporal dynamic (Figure 5-S4 and 5-S5). Finally, we identified genes in each of the five sea star expression clusters with orthologs in each of the planaria and hydra clusters. Using this approach, we find statistically significant overlaps between genes differentially expressed early in all three datasets as well as genes in the posterior-specific sea star cluster with clusters indicating fragment specificity in each of the other organisms. In the following sections we describe how this allowed us to identify not only broad groupings of conserved expression patterns but also specific orthologs similarly expressed across regeneration in these metazoans.

### Early features of the regenerative response are deeply conserved

By analyzing the kinetics of orthologous gene activity in WBR, we find the strongest correlation among genes that are differentially expressed early in each dataset. That is, a significant number of orthologs are up-regulated at early regenerative stages in both the sea star and planaria, as well as the sea star and hydra datasets (hypergeometric p = 4.5 x 10^−3^ and p = 8.8 x 10^−9^, respectively; Figure 5-S4 and 5-S5). This set of genes is enriched for GO terms that include “cilium”, “calcium transport”, and “signaling”. Similarly, we also found a significant number of orthologs are down-regulated in response to bisection in both sea star and planaria (hypergeometric p = 3.3 x 10^−4^). These orthologs are enriched for GO terms such as “ncRNA processing” and “ribosome”, suggesting that early repression of the energetically expensive process of ribosome biogenesis is a fundamental element of WBR.

Two intracellular signaling pathways, Ca^2+^ mobilization and MAPK signaling, have been broadly implicated in wound response [50–54] and are found to be up-regulated early in bipinnaria regeneration. Recent proteomic data indicate that calcium signaling is involved in the anterior regeneration in planaria [55]. MAPK signaling, through both ERK and JNK pathways, is important in neoblast control and blastema differentiation in planaria [56, 57] and JNK signalling has been specifically linked with restoration of proper axial patterning in planaria by re-activation of appropriate WNT signaling [58]. Studies in hydra have similarly demonstrated that wound-responsive MAPK signaling is necessary for early specification of the head organizer, and thus functional regeneration. Early MAPK signaling may thus be shared feature of highly regenerative organisms [59].

The genes upregulated early in regeneration are also enriched for cilium-associated functions. The activation of these genes (*e.g. Ccdc11*, *Rsph3*, *Iqcd*, and *Iqub*; Figure 6A) indicates that, in all three models, cilia play a central role in early regeneration. While this feature has not been reported in either planaria or hydra, a role for cilia in wound response and regeneration has been observed in mammals [45], zebrafish [47], and a related cnidarian (*Nematostella vectensis*) [46].

**Figure 6.**
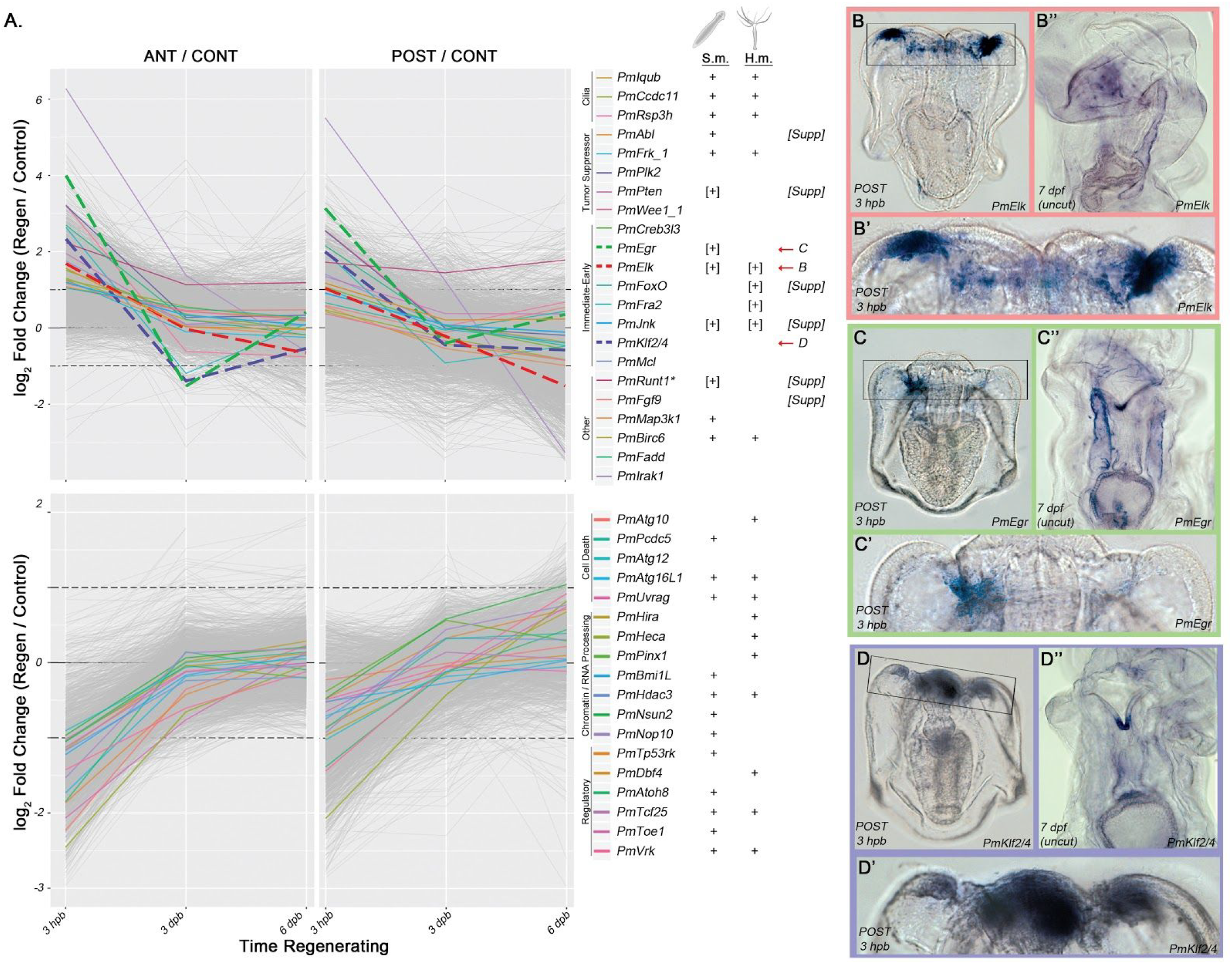
Evolutionarily conserved early regeneration response. (A) These plots show sea star gene log fold change values for genes differentially expressed early in both anterior and posterior regenerating fragments compared with non-bisected sibling control larvae. Genes up-regulated in both fragments (top row) correspond to cluster I and genes down-regulated in both fragments (bottom row) correspond to cluster II. All genes assigned to each cluster are plotted in grey. Several genes, either referenced in the text or representative of functions considered, are indicated with colored lines. Next to the key for each gene is an indication (+) of whether an ortholog for that gene was found in an analogous cluster in either the planaria (*S.m.*) or hydra (*H.m.*) datasets. Indicators in brackets (e.g. “[+]”) are those was no overlapping ortholog identified by our analyses, but genes with the same name were implicated by published datasets. Genes plotted with dashed lines are shown by *in situ* (right). The expression patterns of (B) *Elk* (C) *Egr* and (D) *Klf2/4* are shown. (B’-D’) are magnifications of the wound site shown in the boxed regions in panels (B-D). Expression patterns in uncut larva are also shown (B’’-D’’).

The conserved set of early activated genes also includes several key regulatory genes including orthologs of several tumor suppressor genes (*i.e. Abl*, *Menin*, *Frk, Pten*, *Rbbp6L, Plk2,* and *Wee1*; Figure 6A). Several of these are also upregulated early in other regeneration models [60, 61]; these findings present an additional context in which the tumor suppressor genes show activity during regeneration. In regenerating sea star larvae, normal cell proliferation ceases prior to the emergence of the distinct wound-proximal proliferation (Figure 3). The coincident activation of of tumor suppressor genes and down-regulation of ribosome biogenesis genes may be associated with this response. There is also an early signature of general cell cycle arrest in the hydra transcriptome [5]. While planarian neoblasts continue to proliferate at sites distal to the injury even during blastemal proliferation, inactivation of planarian *PTEN* gene homologs resulted in defective regeneration due to neoblast hyperproliferation [62]. These results indicate that a common early feature of WBR in these systems is the modulation of regulators of cell proliferation.

In addition to cell proliferation, these analyses suggest that cell death is tightly regulated early in regeneration. Genes associated with regulating cell death pathways are another example of conserved differential expression early in these models. Notably, at least seven genes in the autophagy pathway are down-regulated in regenerating sea star larvae, planaria, and hydra (i.e. *Atg16L1, Atg12*, *Atg10, Atg14*, and *Uvrag*; Figure 6A). This is consistent with findings in hydra that suggest autophagic cell death is repressed during regeneration [63]. Conversely, as autophagy is downregulated in sea star larvae, genes that modulate apoptotic cell death are activated (e.g. *Fadd*, *Birc6*, and *Ulk1*). Apoptotic cell death is necessary for increased i-cell proliferation in hydra [18] and, in planarian regeneration, has been implicated in tissue remodeling and neoblast proliferation [64, 65]. Despite these early transcriptional changes, an increased number of TUNEL^+^ cells is not apparent until much later in bipinnaria regeneration (6 dpb; Figure 4). Therefore this modulation in cell death may be pathway specific (i.e. autophagy vs apoptosis) or otherwise undetected by our TUNEL assay. Alternatively, these transcriptional changes may be involved in establishing an appropriate balance between cell death and cell proliferation during this early phase.

Finally, we identified a suite of immediate early genes that are activated in all three animals. In regenerating sea star larvae, we find rapid, significant up-regulation of *Jnk*, *Elk*, *Egr*, *Klf2/4, Mcl*, *Creb3l3*, *Fra2*, and *FoxO* (Figure 6A). For example, *Egr* is one of the most strongly up-regulated genes in both anterior and posterior regenerating sea stars (Figure 6C), while in planarian regeneration *EGR* is one of the earliest and strongest wound-proximal genes induced during planarian regeneration [10]. The conserved early down-regulation of the *Egr* repressor *Toe1* in both sea stars and planaria suggests these genes are parts of concerted early response in these contexts. Several of these early activation factors are also known to be regulated by MAPK signaling pathways in other systems [66]. For example, in the sea urchin *Strongylocentrotus purpuratus*, *SpElk* is a target of MAPK signaling (ERK) and regulates both *SpRunt1* and *SpEgr* expression during embryogenesis [67]. In planaria, MAPK signaling (*Jnk*) activates *Runt1* and *Egr* following wounding [65]. *Jnk* signaling in hydra has been shown to regulate *FoxO* expression [68], which is an important regulator of hydra i-cells [69].

These overlapping sets of genes differentially expressed early reflect a common response to the bisection insult. This suggests that these gene orthologs define key shared characteristics between highly regenerative species in a specific response to injury that permits the regeneration program.

### Genes underlying deeply conserved early response are dramatically upregulated in the sea star wound site

We additionally chose a subset of these genes to examine their spatial localization during regeneration. *Elk* and *Egr* are both normally expressed in coelomic pouch epithelium (Figure 6B’’-C’’), but by 3 hpb they are also strongly expressed in the sites of wound closure (Figure 6B’-C’, Figure 6-S1A-B). *Fgf9* expression is also localized to wound sites during early regeneration (Figure 6-S1 F). Although neither *Ets* nor *Erg* were siginificantly differentially expressed by RNA-seq or nanostring, we examined their expression given their known expression in sea star mesenchyme [70]. We find that both are localized to wound sites during early regeneration (Figure 6-S1 D-E), suggesting an early role for mesenchymal cells, although not necessarily due to a transcriptional change. *Klf2/4* is normally expressed strongly in the mouth and foregut and after bisection is strongly upregulated in wound-proximal foregut (Figure 6D’ and Figure 6-S1C). Conversely, *FoxO, Jnk,* and *Runt* are expressed in the tip of the foregut proximal to the wound site, but not in the wound itself (Figure 6-S1 G-I). The tumor suppressor genes *Abl* and *Pten* are expressed broadly around the wound during early regeneration (Figure 6-S1 J-K). This spatial expression therefore shows that the set of gene homologs with early regenerative response among these deeply divergent animals are expressed in the early wound region of the sea star larva.

### Axis respecification precedes wound-proximal proliferation

Restoration of normal gene expression levels along the bisected AP axis must be a central component of regeneration. Gene expression domains for components of the GRN that controls early axial patterning in sea star embryo have been well defined. The Wnt pathway, for example, has well characterized functions in specifying the embryonic AP axis [49, 70]. Anterior ectoderm domains required for the development of the larval nervous system have also been delineated [71–73]. This enables us to analyse the expression of these genes during regeneration. And indeed, analysis of genes within the two expression clusters differentially expressed in regenerating anterior and posterior larval fragments (Clusters III and IV; Figure 5) demonstrates that embryonic axis patterning genes are expressed during AP axis restoration.

When examining these clusters, it should be noted that although genes in these clusters appear to be rapidly down-regulated following bisection, because transcript levels were normalized to those in whole larvae, this phenomenon is actually a result of removing cells and tissues in the other half of the larva. For example, genes normally expressed in anterior larval domains (e.g. *Frizz5/8* and *FoxQ2*) initially appear to be down-regulated in posterior fragments relative to uncut larvae but are unaffected in anterior fragments (solid lines, Figure 7; Cluster III, Figure 5). Correspondingly, genes that are typically expressed in the posterior domain (e.g. *Frizz9/10*, *Wnt16*, and *Nk1*) are absent in anterior fragments but unaffected in posterior fragments (dashed lines, Figure 7; Cluster IV, Figure 5). For several genes in each of these clusters expression levels recover to pre-bisection levels within 6 days. Notably, however, this processes appears to be delayed within the regenerating anterior fragments relative to the posterior fragments (Figure 7).

**Figure 7.**
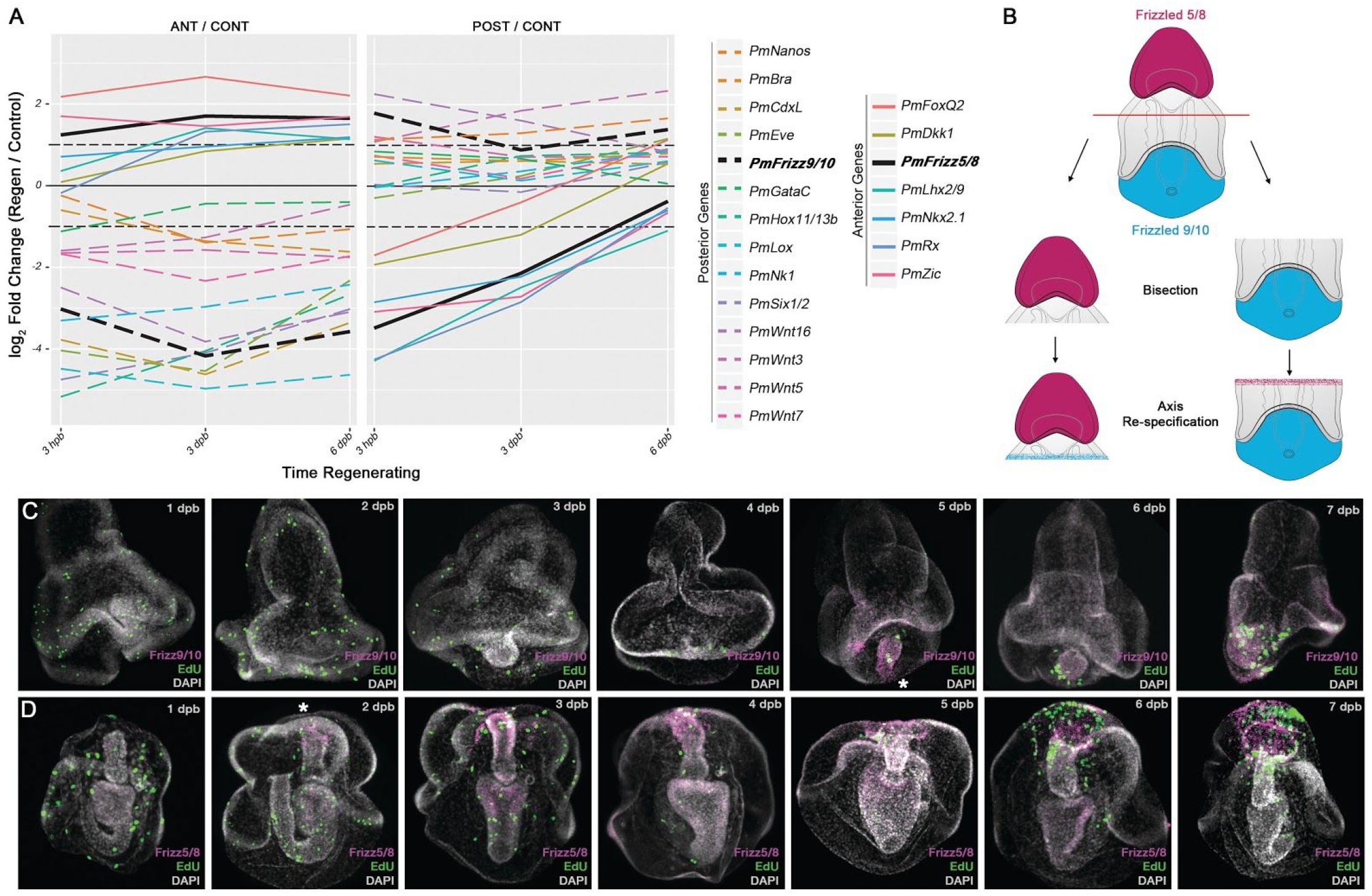
Fragment-specific recovery of appropriate anterior-posterior gene expression. (A) The expression of genes asymmetrically expressed in either anterior (ANT; solid lines, Cluster III) or posterior (POST; dashed lines, Cluster IV) sea star larval territories was examined at 3 hours post bisection (hpb), 3 days post bisection (dpb) and 6 dpb. The log fold change values for each gene in regenerating anterior or posterior fragments compared with non-bisected sibling control larvae is reported for each fragment (ANT/CONT and POST/CONT, respectively) over the regenerating time course sampled. Black lines show the detected expression of *Frizz5/8* and *Frizz9/10*. (B) Model for recover of genes asymmetrically expressed along the anterior-posterior axis, with *Frizz9/10* (blue) and *Frizz5/8* (maroon) provided as examples. (C) Whole mount fluorescent *in situ* hybridization illustrating the re-activation of *Frizz9/10* (magenta) in the posterior aspect of regenerating anterior fragments beginning at 5 dpb and preceding the concentration of proliferating EdU^+^ cells (green) near the wound site. (D) Re-activation of *Frizz5/8* (magenta) in the anterior aspect of regenerating posterior fragments beginning at 2 dpb and preceding the concentration of proliferating EdU^+^ cells near the wound site.

To characterize more fully, the reestablishment of axial patterning during regeneration we examined the spatial expression of two Wnt pathway receptor genes: *Frizz5/8* (normally expressed in the anterior), and *Frizz9/10* (localized in the posterior). In the anterior regenerating fragments, *Frizz9/10* transcripts are undetectable following bisection (immediately after the posterior halves were removed). However, by 5 dpb *Frizz9/10* transcripts are evident in the the newly developed posterior domain (Figure 7C). Additionally, we detect the re-expression of *Frizz9/10* before the onset of wound-proximal proliferation. Likewise, *Frizz5/8* is undetectable in regenerating posterior fragments until about 2 dpb when it is seen in the anterior aspect of these fragments (Figure 7D), again before proliferating cells localize to this region. Appropriately localized expression of *Frizz9/10* and *Frizz5/8* persists in regenerating posterior and anterior fragments, respectively (Figure 7-S1B,E). This finding extends to other genes with known roles in embryonic AP axial patterning that are identified in our clusters. For instance, We find similar recapitulation of embryonic expression patterns for e.g, *FoxQ2* (another anterior marker) and *Wnt8* (an additional posterior marker; Figure 7-S1 F-J). Thus embryonic patterning genes are used again during the restoration of the AP axis, and this precedes the initiation of blastemal proliferation.

This pattern mirrors planarian regeneration in which blastema formation and regeneration cannot proceed when axis specification is perturbed [74–76]. Although hydra regeneration does not require blastemal proliferation, interstitial cells proliferate following wounding and this proliferation is initiated by a transient release of Wnt3, a protein implicated in head organizer function [18]. This comparison between animals positioned across the metazoa suggests the important finding that regeneration-associated proliferation requires a resetting of an axial positional program.

### Common regulatory toolkit used for axial respecification

We sought to determine if any of the genes involved in sea star axis respecification during regeneration are conserved among animals. We examined the genes assigned to these fragment-specific clusters (Clusters III and IV) to identify orthologous genes with similar expression trends in the other datasets. We find significant overlaps between the posterior-specific sea star genes (Cluster IV) and asymmetrically expressed genes in both hydra (Cluster 1, Figure 5-S5) and planarian (Cluster 2, Figure 5-S4) datasets. The hydra oral-aboral axis corresponds to the posterior-anterior axes in bilaterians [77]. The RNA-Seq data from hydra were generated using oral regions of the regenerating aboral body stalk [5]. Thus, the signature of late stage up-regulation reflects the recovery of transcripts typically expressed in the head (Cluster 1, Figure 5-S5) and we expect that oral gene expression in hydra would correspond to posterior gene expression in sea stars. These nominally oral-specific genes in hydra in fact do exhibit a significant overlap with the posterior-specific sea star genes (hypergeometric p = 2.7 x 10^−3^). Likewise, genes asymmetrically expressed between anterior and posterior halves in the planaria dataset overlap the posterior-specific sea star genes (hypergeometric p = 1.4 x 10^−2^). In both cases, the overlapping genes include Wnt ligands and receptors (e.g. *Wnt7, Wnt5,* and *Frizz9/10*) and other regulatory genes associated with posterior fates (e.g. *Bra, Hox11/13a,* and *Six1/2*). The observed overlap in asymmetrically expressed genes among these datasets suggests a that a common regulatory toolkit is deployed for axis respecification in each of these models that includes Wnt signaling. The absolute orientation of the axes is not conserved, but this likely reflects developmental usage.

### Temporal dynamics of regeneration induced cell proliferation differ among these animals

The patterns of cellular proliferation is one aspect in which the three models of WBR differ considerably. Sea star larvae and planaria exhibit concerted wound-proximal proliferation that coincides with the final time points sampled here: 6 dpb for sea star larvae and 3 dpb for planaria. Early in planarian regeneration, a global burst of neoblast proliferation is also observed (i.e. within 6 hours post amputation) [9]. No such early increase in proliferation is observed in sea star larvae (Figure 3). While hydra do not rely on a proliferative blastema to resupply cells for regeneration, interstitial stem cells (i-cells) proliferate proximal to the wound within the first 2-4 hours post amputation [18]. This i-cell proliferation follows the early suppression of mitosis that is observed after wounding [5].

In sea star larvae, the genes up-regulated later in regeneration in both the anterior and posterior fragments (Cluster V; Figure 5) are strongly associated with cell proliferation. It is important to note that while overall numbers of proliferation cells are decreasing, the timing of the up-regulation of these genes correlates with the emergence of wound localized proliferation. We compared these genes with orthologs that exhibit similar expression dynamics in the other datasets. None of the expression clusters from planaria or hydra are significantly enriched in orthologs of the sea star proliferation genes. Specifically, very few orthologs are apparent between the later upregulated sea star cluster (Cluster V) and the corresponding gene clusters from planaria and hydra (i.e. planaria Cluster 1 and hydra Cluster 3; Figure 5-S4,5). Instead, there is a strong, though not statistically significant, overlap between the genes up-regulated late in sea star and those up-regulated early in planaria (e.g. Cluster 3, Figure S3) and hydra (e.g. Cluster 1, Figure S4). Many of these shared genes are associated with cycling cells (e.g. DNA polymerase subunits, MCM genes, structural maintenance of chromosomes [SMC] genes, *Orc3*, *Rrm1*, *Plk*, and *Ttk*). These data suggest the intriguing hypothesis that wound-proximal proliferation in sea star larvae is more similar to early bursts of cell proliferation than the later blastemal proliferation observed in planarian regeneration.

### Regeneration induces coincident expression of normally tissue restricted proliferation-associated genes

We examined the expression patterns of proliferation-associated genes during regeneration (i.e. Cluster V). *Mcm2*, *Runt1*, *GliA*, and *Dach* are all expressed in the anterior region of regenerating posterior fragments, coincident with the wound-proximal proliferation (Figure 8 B-E). Each gene is expressed in multiple distinct tissues, including the anterior foregut, anterior epithelium, coelomic epithelium, and gut (Figure 8B-E). Notably, however, during embryonic and larval stages these genes exhibit non-overlapping expression patterns. For example *Mcm2* is expressed in the ciliary band and foregut, *Runt1* is expressed in the mouth, mid-gut and hindgut, *GliA* is strongly associated with the developing coelomic epithelium, and *Dach* is expressed throughout the gut and in ciliary band epithelium (Figure 8-S1). These results indicate that a suite of genes that function in cell proliferation and are normally expressed in diverse tissues are re-deployed during regeneration and are co-expressed in the proliferating blastema.

**Figure 8.**
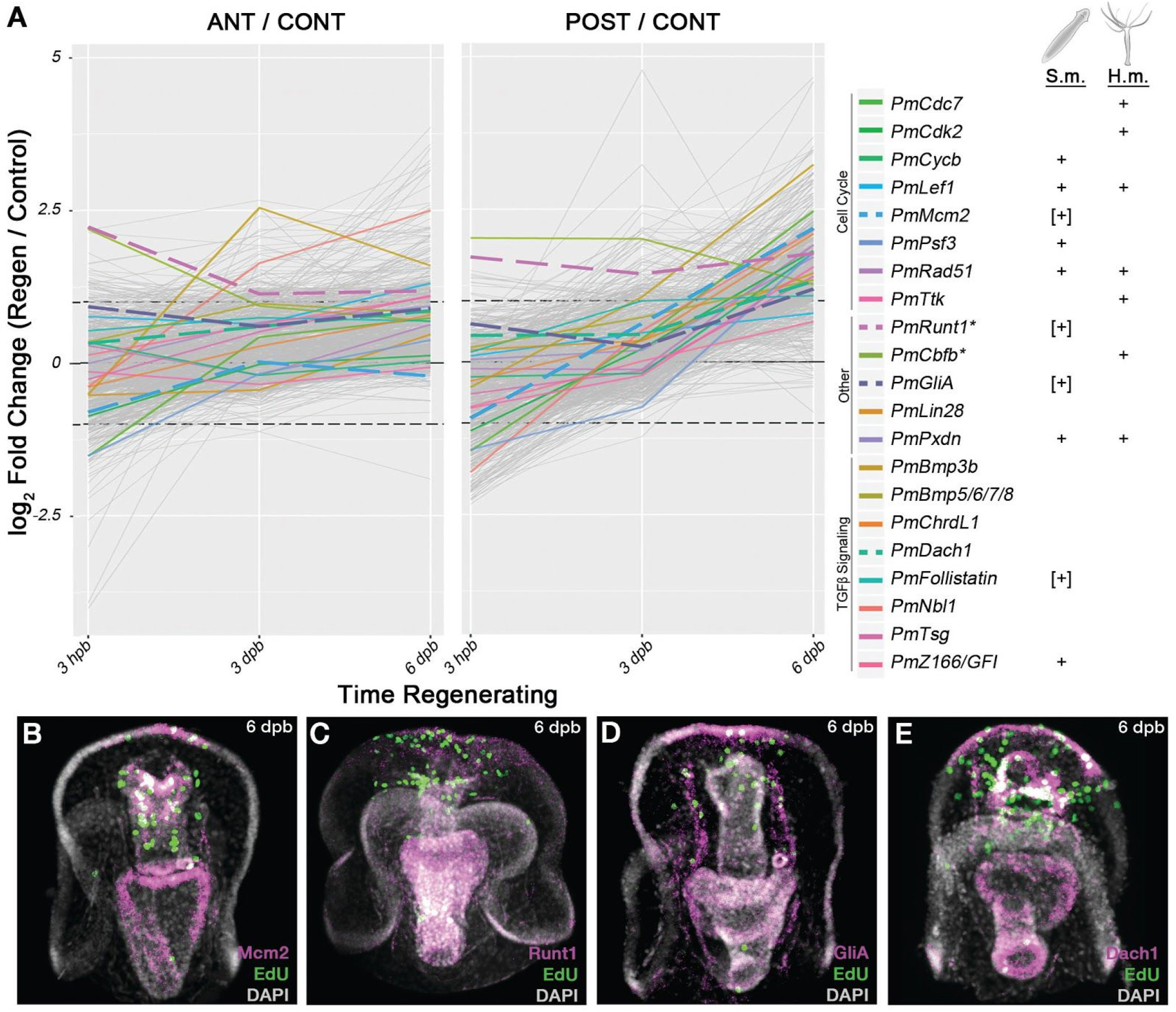
Conserved proliferation-associated genes. (A) These data show sea star log fold change values for genes differentially expressed at later stages in regenerating fragments compared with non-bisected sibling control larvae (i.e. sea star cluster V). All genes assigned to cluster V are plotted in grey. Several genes, either referenced in the text or representative of functions considered, are indicated with colored lines. Next to the key for each gene is an indication (i.e. “+”) of whether an ortholog for that gene was found in an analogous cluster in either the planaria (*S.m.*) or hydra (*H.m.*) datasets. Indicators in brackets (e.g. “[+]”) are those where no overlapping ortholog was identified by our analyses, but genes with the same name were implicated by published datasets. Genes plotted with dashed lines are shown by fluorescent *in situ* hybridization (below). *Mcm2* (B), *Runt1* (C), *GliA* (D) and *Dach1* (E) are all expressed in the anterior aspects of regenerating fragments at 6 dpb. In many cases, the expression of these genes is coincident with an EdU^+^ cell, suggesting that these genes are expressed, at least in part, in proliferating cells.

## CONCLUSION

While the capacity for larval sea stars to undergo WBR has been appreciated for over two decades, there has not yet been a systematic characterization of the cellular and molecular processes involved. In the present study we demonstrate that larval sea stars exhibit many stereotypical characteristics found in other models of WBR. This is a striking finding because sea stars are Deuterostome animals and very distantly related to the other species considered here. Through our transcriptome analyses, we detect an early wound-response phase involving significant alterations in the expression of stress response genes, genes involved in signaling pathways (including MAPK, Ca^2+^) and a broad shut-down of energetically expensive anabolic processes (e.g. ribosome biogenesis). The first few days following bisection are marked by a global decrease in the number and distribution of cycling cells compared to what is typically observed in growing, non-bisected larvae. This precedes, the re-establishment of developmental axes, specifically the AP axis ablated by bisection. Re-patterning of the AP axis is observed both through *in situ* hybridization as well as transcriptome measurements. These observations are facilitated by our extensive prior knowledge of sea star developmental patterning programs and, indeed, genes described by the developmental gene regulatory network are enriched in these clusters. Notably, through both our transcriptome and *in situ* experiments we observe that axis respecification occurs prior to the onset of wound-proximal cell proliferation, which is the last phase assayed in the present study. This is the first description of concerted, wound-proximal cell proliferation in regenerating sea star larvae. Given that this wound-facing region in both regenerating fragments is the primordium from which larval tissues regenerate, we define this proliferative zone as the regeneration blastema. In this study we have only monitored the first half of the regeneration process up until the emergence of this blastema. Complete regeneration in these larvae takes a total of 10-14 days [42, 43].

In this work we sought to leverage the power of comparing regeneration in a variety of contexts to identify common features. For example, we clustered gene expression levels to identify genes similarly differentially expressed in both anterior and posterior regenerating sea star larval fragments. These patterns were then used as a basis for comparison to planaria and hydra regeneration datasets. In the present study we compared regeneration in species that last shared a common ancestor approximately 580 million years ago, at the base of the metazoa. This is the broadest direct comparison of regeneration yet described, encompassing three of the major groupings of animals (Deuterostome, Protostome, and basally branching Eumetazoa). We find evidence for conservation of both broad functional classes as well as specific orthologs involved with the regenerative process among these animals. Although there are examples in our data of genes with divergent expression patterns in these various animals, we focus our attention on those that are are conserved as these have the greatest potential to inform our goal of identifying common features of highly potent regeneration. Indeed, the genes found to be in common are eukaryotically conserved orthologs with deeply conserved functions in core cellular processes that are required in many regenerative contexts (e.g. cellular proliferation, apoptosis). The significance of our finding here is not that we detect such genes, but that we find evidence for conservation of temporal expression in many of these processes. As with any EvoDevo study, it is difficult to absolutely distinguish between a genuine homology of these regenerative programs, rather than independent convergence of multiple critical pathways. We think, however, that the extensive conserved temporal ordering of these processes (e.g. axial respecification prior to cell proliferation) points toward conservation. These commonalities are summarized in Figure 9. The most remarkable signature of shared genes and processes is among genes both up-and down-regulated early. We are potentially most empowered to detect such an overlap among early genes as temporal synchrony between the models likely diverges later in the time course. Nonetheless early changes to Ca^2+^ and MAPK signaling pathways, upregulation of ciliogenesis genes, upregulation of tumor suppressor genes, downregulation of autophagy genes, and activation of a suite of immediate early genes are common aspects of regeneration in these models. There is also a set of conserved genes that we hypothesize are commonly involved in axial respecification, most notably genes in the WNT signaling pathway. Importantly, axis respecification occurs prior to regeneration associated proliferation across these species. In contrast to these commonalities, we show that the temporal profiles of gene expression underlying the proliferative response are different.

**Figure 9.**
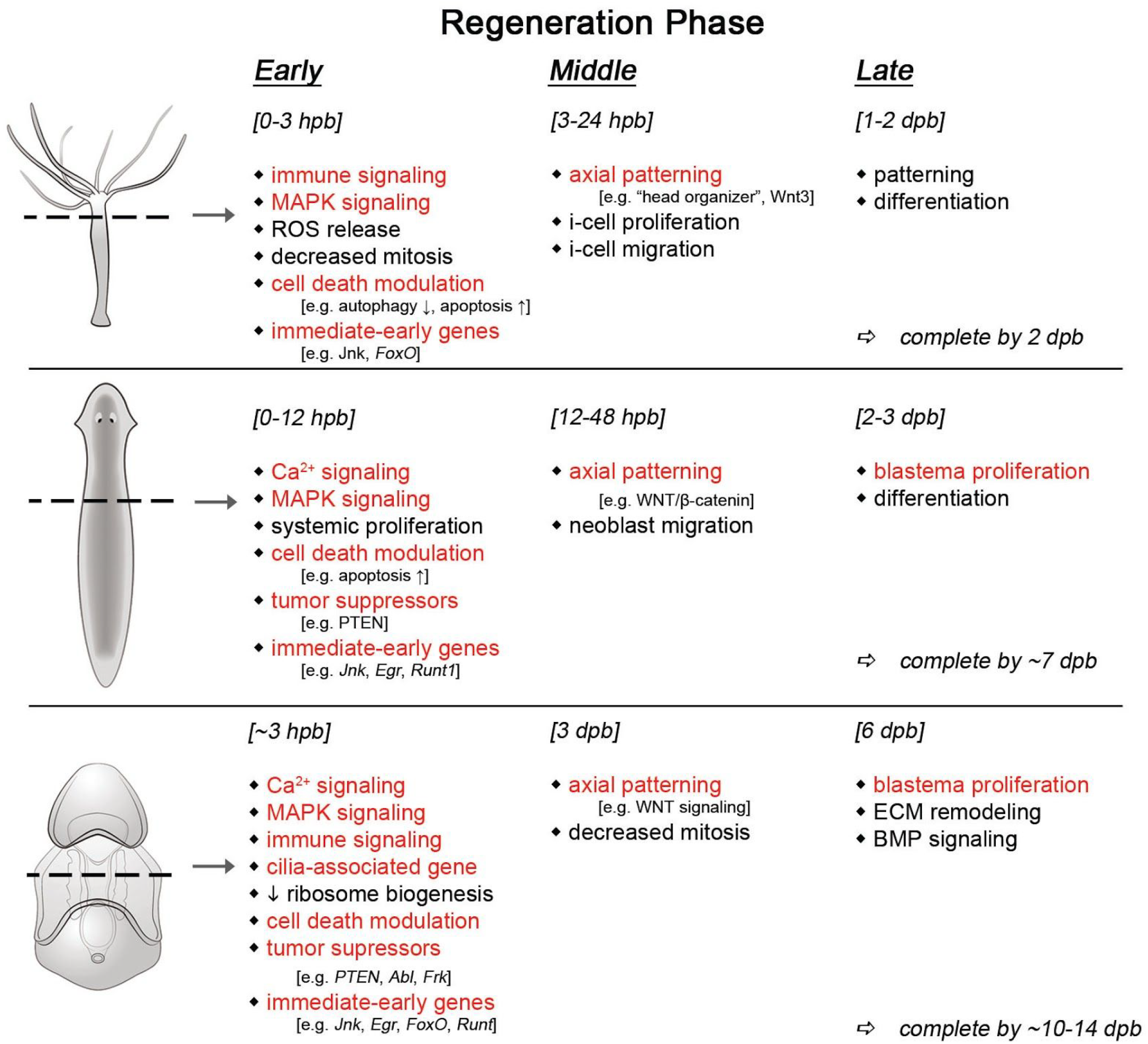
Summary of similarities between WBR models. The reported features of regeneration at early, middle, and late stages of regeneration, with respect to the datasets considered in this study, are indicated. Features detected in the sea star model in our study that are shared with the other two models are highlighted in red. Some aspects are considered in common based on shared gene expression (e.g. MAPK signaling) whereas others are based on cytological observations (e.g. blastema proliferation).

These commonalities between highly diverged WBR models highlights a deep conservation in regeneration mechanisms among the metazoa. This work also underscores the power of comparative inquiries in identifying the core components of the regenerative response and, potentially, how these components are altered in non-regenerative species.

## METHODS

### Animals and Regeneration Paradigm

Adult *Patiria miniata* were obtained from the southern coast of California, USA (Pete Halmay or Marinus Scientific) and were used to initiate embryo cultures as previously described [78]. *P. miniata* embryos were cultured in artificial seawater at 16 °C and fed *Rhodomonas les* ad libitum every 2 days along with fresh artificial seawater beginning at 4 days post fertilization (dpf). All studies of regenerating larvae were conducted with larval cultures beginning at 7 dpf at which point the larvae were manually bisected stereotypically through the foregut, midway along the transverse anterior-posterior axis (Figure 1B), with a #11 sterile scalpel. Resulting anterior and posterior fragments, as well as control (uncut) larvae, were then transferred to separate 35 mm polystyrene dishes at a density of no more than 50 larval fragments per ml of artificial seawater and cultured for the time indicated.

### Whole-Mount Staining and Staining Larval Sections Procedures

*P. miniata* larvae or regenerating larval fragments, grown for the times indicated, were fixed in a solution of 4% paraformaldehyde in MOPS-fix buffer (0.1M MOPS pH 7.5, 2mM MgSO_4_, 1mM EGTA, and 800 mM NaCl) for 90 minutes at 25 °C and transferred to a solution of 70% ethanol for long term storage at −20 °C. *In situ* hybridization experiments were performed as previously described [71, 79] using digoxigenin-labeled antisense RNA probes. Labelling and detection of proliferating cells in *P. miniata* larvae was performed using the Click-it Plus EdU 488 Imaging Kit (Life Technologies), with the following modifications. Larvae were incubated in a 10 μM solution of EdU for 6 hours immediately prior to fixation in a solution of 4% paraformaldehyde in phosphate buffered saline (PBS). Larvae were fixed for a minimum of 90 minutes at 25 °C and subsequently transferred to a solution of 70% ethanol for storage at −20 °C. For detection of EdU incorporation, labeled embryos were transferred to a solution of PBS and the detection was performed following the manufacturer’s protocol.

For detection of *in situ* and EdU staining in the same specimen, EdU labelled larvae were fixed and hybridized with digoxigenin labelled riboprobes, as described. Detection was performed using anti-digoxigenin POD-conjugate antibody (Sigma) and a tyramide signal amplification (Perkin Elmer). Following signal deposition, larvae were washed in PBS and EdU was detected as described.

For BrdU pulse-chase experiments, larvae were labeled with 50 μg/ml solution of BrdU (Sigma B5002) for 6 hours after which they were washed and placed in fresh seawater. Following fixation, larvae were denatured in 2 M HCl and 200mM NaCl for 30 minutes at 37 °C. The denaturant was neutralized in 0.1 M Borate buffer (pH 8.5), followed by blocking in PBS with 2% BSA and 0.1% Tween-20. The anti-BrdU antibody (Sigma B2531) was diluted 1:100 in blocking buffer incubated for 1 hour. The larvae were then washed in PBS with 0.5% Tween-20 and incubated in a 1:500 dilution of anti-mouse Alexa 568 (Life Technologies) for 1 hour. Following additional PBS washes EdU detection was performed as described.

For TUNEL staining, animals were fixed in 4% paraformaldehyde in 0.01 M phosphate buffer (pH 7.4, 1007 mOsm) for 24 h at 4 °C. After fixation the embryos were incubated in 0.1M glycine in phosphate buffer with 0.1% Tween 20 for 1 hour to quench residual autofluorescence. The tissue were permeabilized in 0.5% Triton X-100 for 30 min and by Proteinase K digestion (8 μg/ml, 10 min at room temperature). Cells undergoing programmed cell death were identified using the Fluorescent FragEL™DNA Fragmentation Detection Kit (Calbiochem) as per the manufacturer’s protocol. Images of whole mount specimens were taken using the Zeiss LSM 880 scanning laser confocal microscope. Maximum intensity Z-projections and automatic cell counting were generated in the Fiji image processing software.

At least two independent biological replicate experiments were performed for each *in situ*, EdU staining, or TUNEL staining experiment, examining the pattern of at least 10 specimens per replicate. For quantitation of EdU and TUNEL images, Z-projections were generated and counted in ImageJ. Images were converted to 16-bit prior to thresholding. For images of anterior larval segments, a 0.4% threshold was used, and for images of uncut larvae and posterior segments, a 1% threshold was used. Each image was then converted to a binary mask shed. Using the Watershed tool, larger objects were segmented into individual cells. To segment each image into three sections (wound, middle, distal), each image was divided into three equal portions. To quantify the number of EdU^+^ cells in each section, the Analyze Particles tool was used. For uncut larvae the size parameters used was 5-300μm^2^ and for regenerating larvae, 20-300μm^2^. Statistical analysis of count data was performed using the estimation stats website [80] and, in all cases, used the 0 dpb as a shared control sample and reported p-values are based on nonparametric Mann-Whitney U tests.

For histology, larvae were fixed in 4% paraformaldehyde in 0.01 M phosphate buffer (pH 7.4, 1007 mOsm). After fixation, the specimens were rinsed in the same buffer and postfixed in 1% OsO_4_ for 1 hour. The samples were dehydrated in a graded series of ethanol and propylene oxide and embedded in the Araldite epoxy resin. Sections were cut with glass knives on Ultracut E (Reichert, Vienna, Austria). The serial semi-thin (1 μm) sections were collected on gelatin-coated slides, stained with 1% toluidine blue in 1% aqueous sodium borate and mounted in DPX (Fluka). The sections were viewed and photographed with a Leica DMI 4000B microscope equipped with a Leica DFC 420C camera.

### RNA-seq, Read Mapping, and Transcriptome Assembly

For transcriptome measurements, larvae were grown and bisected as described in results. RNA was collected from pools of approximately 300 sibling individuals of regenerating anterior fragments, regenerating posterior fragments, as well as uncut control larvae. Two biological replicate samples were prepared for each timepoint for a total of 18 samples. RNA was extracted using the GenElute Mammalian Total RNA Kit (Sigma-Aldrich). Illumina TruSeq library preparation and HiSeq 2500 50 bp SR sequencing were performed (USC Epigenome Center).

RNA-seq reads were trimmed of residual adapter sequences and low quality bases (Trimmomatic v0.32, [81]). High-quality reads were mapped to the *P. miniata* v1.0 genome assembly (Tophat v2.0.12, [82]) and, in total, 422.9 M uniquely mapping reads were recovered from the 18 samples at an average depth of 23.5 M reads per sample. Uniquely mapping reads were assembled into transcripts using Cufflinks [83] and the MAKER2 based gene predictions hosted at Echinobase were used to guide transcript assembly. Reads uniquely mapping to a gene (locus) from this Cufflinks transcriptome assembly were counted (HTSeq-count v0.6.1p1, [84]). Read counts were normalized and genes detected with more than 3 reads per million, corresponding to 50-120 uniquely mapping reads depending on the sample, in at least two samples were retained for further analyses, corresponding to 31,798 expressed genes. Raw and processed sequencing reads have been deposited into the NCBI Gene Expression Omnibus (GSE97230) and analysis scripts are available upon request.

### Gene Ontology term annotation and Ortholog identification

The newly assembled sea star genes were annotated in three ways: by identifying the reciprocal best BLAST hit (rBBH) between the sea star transcript and either sea urchin or mouse genes and using Blast2GO. 9,027 (28.4%) loci have an rBBH match to a sea urchin protein, 7,212 (22.7%) loci have an rBBH match to a mouse gene, and 9,617 (30.2%) assembled loci were annotated using Blast2GO. GO terms for each sea urchin and mouse gene were assigned to their respective rBBH match in the sea star set and these were used for enrichment analyses. Overall the results based on all three annotation methods are highly similar (Figure 3B and Figure S2). Reciprocal best BLAST hits (rBBH) were also used to identify putative orthologs between the sea star genes and the planaria and hydra transcripts. We found 5,220 *S. mediterranea* transcripts and 6,091 *H. magnipapillata* transcripts with an rBBH match to a sea star transcript.

### Differential Expression Testing and Hierarchical Clustering

Expression levels in biological replicate samples are highly correlated (pearson correlation coefficient = 0.985). Regenerating fragments were compared to age-matched sibling uncut control larvae and differential expression was assessed using a generalized linear model quasi-likelihood F test (edgeR, [85, 86]), controlling for sample batch. Differentially expressed genes (DEG) were defined as those changes detected below a p-value of 0.05 and with a fold change greater than 2-fold in either direction. Using these criteria there are 9,211 total DEG in at least one regenerating fragment compared to the control larvae and at least one of the timepoints sampled, which represents 28.97% of all of the expressed genes detected.

The fold-change values for all 9,211 DEG relative to control larvae were clustered by first computing the euclidean distance matrix and then these values were then clustered using the “ward.D2” method provided as part of the R hclust function. The optimum number of clusters was determined by cutting the resultant dendrogram at various heights and empirically determining at which height the number of clusters began to plateau (h=42). The result was 8 distinct clusters. However, we noted that several clusters shared similar overall patterns (Figure 5-S1). As the similar clusters shared very similar GO enrichments and expression patterns over the time course, we further grouped these into the final 5 clusters reported in the text. The grouping of clusters did not alter the enrichment of GO terms or our other downstream analyses (Figure 5-S3).

For the planaria and hydra regeneration datasets, data was obtained from supplemental tables associated with each publication. The planarian data were reported as normalized read counts for the 15,422 transcripts detected. These counts were log_2_-transformed and then scaled to z-scores, or the number of standard deviations from the mean value for each transcript, and only those transcripts considered differentially expressed as reported by the authors were considered. This resulted in 7,975 transcripts that were then clustered in the same way as described above for the sea star transcripts. The hydra data were reported as binned z-scores for the 28,138 transcripts detected corresponding to lower, mid, and upper third of expression range for each transcript. We only clustered transcript values for which a positive reciprocal match was detected, leaving 5,779 transcripts for our analyses. The euclidean distance matrix was calculated, as with the other datasets, but to accommodate the binned nature of these data the hierarchical clustering was performed using the “average” method provided with the hclust R function. A fine-grained resolution of common gene expression dynamics across these species is not warranted without more closely aligning experimental designs, including sampling time points and normalization strategies. Therefore, for each of these datasets we sought very broad cluster classifications such that assigned genes are either either up-regulated early and down later or vice versa in their respective time course. The result is three clusters each for the *S. mediterranea* and *H. magnipapillata* datasets (Figures 5-S4 and S5).

### Nanostring nCounter Assay Analysis

A custom Nanostring nCounter codeset was designed, available upon request, consisting of 114 total probes - 8 negative control, 6 postitive control, 11 housekeeping control, and 89 gene-of-interest probes. RNA was prepared from similarly staged larvae and hybridized to the codeset as directed by the manufacturer. The nCounter DA71 digital analyzer output files were collected and further analysis was performed using the NanoStringDiff R package [87]. Briefly, background signal was defined for each sample as the mean plus two standard deviations of the negative control probes and assigned as the negative control normalization factor parameter. The geometric mean of signals for each sample from positive control probes and housekeeping probes were used to calculate a positive control and housekeeping scaling vectors for each sample. Differential expression between regenerating fragments and control, uncut larvae was determined using a generalized linear model likelihood ratio test (p < 0.05). Probes that failed to express above background levels were omitted from further analyses. Finally, heatmaps of fold-change calculated based on Nanostring measurements were plotted for genes assigned to groups based on RNA-seq cluster identities. Genes with similar general expression dynamics (e.g. up early in both fragments, down early in both fragments, etc) in both RNA-seq and Nanostring experiments were detected.

## LIST OF ABBREVIATIONS

WBR: whole-body regeneration
GRN: gene regulatory network
DEG: differentially expressed gene
GO: gene ontology
hpb: hours post bisection
dpb: days post bisection
WMISH: whole mount in situ hybridization
ANT: anterior
POST: posterior
CONT: control
AP: anterior-posterior
rBBH: reciprocal best blast hit
S.m.: *S. mediterranea*
H.m.: H. magnipapillata
N.S..: not significant

## DECLARATIONS

### Ethics approval and consent to participate

Not applicable.

### Consent for publication

Not applicable.

### Availability of data and material

Datasets generated during the current study are available from NCBI GEO (GSE97230). Planarian transcriptome data have been published [4]. Hydra transcriptome data have been published [5] and transcriptome sequences are available from Thomas Holstein, upon request.

### Authors’ contributions

VH and GC conceived of and designed experiments. GC, AW, OZ, and JP carried out experiments. GC, AW, and OZ analyzed data and GC wrote the manuscript. VH was instrumental in the preparation of the manuscript and interpretation of the results. All authors read and approved of the final manuscript.

## Acknowledgements

The authors thank Thomas Holstein for sharing the hydra transcriptome assembly; Sara J Cary Illustrations and Photography for providing illustrations of bipinnaria, planaria, and hydra; Katherine Buckley, Minyan Zheng, and William Hattleberg for helpful feedback during the preparation of the manuscript.

